# Respiratory viral infection is associated with increased *Pseudomonas* abundance in cystic fibrosis airways

**DOI:** 10.1101/2025.07.08.663034

**Authors:** Yasmin Hilliam, Catherine R Armbruster, Samar E Atteih, Glenn J Rapsinski, John Moore, Junu Koirala, Leah Krainz, Jordan Gaston, John Williams, Vaughn S Cooper, Stella E Lee, Jennifer M Bomberger

## Abstract

Acute respiratory viral infections are an important driver of morbidity and mortality in people with chronic lung disease and are frequently associated with pulmonary exacerbations and a transition from intermittent to chronic bacterial infection of the airways. Chronic *Pseudomonas aeruginosa* infections are associated with worsened lung function, poor outcomes, and increased hospital visits. We sought to improve understanding of the effects of respiratory viral infections and host immune response on the resident bacterial community of the airways, using cystic fibrosis as a model. We performed an observational longitudinal study of 38 adults with CF and collected sinus and sputum samples at 6-month intervals from 2017 – 2021. We performed 16S rRNA amplicon sequencing to characterize the airway microbiota, real-time RT-PCR for viral infection detection, and cytokine quantification. We observed viral positivity rates of 19% and 14% in sinus and sputum samples, respectively. Human rhinovirus was the most frequently observed viral pathogen in both sinus and sputum samples. We measured a significant perturbance of the bacterial community during viral infection that did not return to baseline following resolution of the viral infection. This perturbation was driven by a significant increase in *Pseudomonas* relative abundance during viral infection. Furthermore, we found significant associations with increased *Pseudomonas* relative abundance for several pro-inflammatory and antiviral cytokines, including interleukin (IL)-2, IL-8, and interferon (IFN)-*λ*1. These findings indicate an important role for respiratory viral infections and the host immune response in the development and maintenance of chronic *Pseudomonas* infections in the context of CF airway disease and broadly expand our understanding of viral-bacterial coinfection of the airways.

**IMPORTANCE:** Respiratory infections are a leading cause of morbidity and mortality worldwide, and co-infections are associated with worsened disease outcomes. In viral-bacterial co-infection, clinical and mechanistic studies show that a preceding acute respiratory viral infection promotes the establishment and exacerbation of bacterial infections, leading to increased morbidity. Although people with cystic fibrosis do not experience more frequent acute respiratory viral infections, their outcomes are worse, with prolonged symptoms and hospitalizations. When examining how acute viral infections shape the microbiota in the respiratory tract of pwCF, we observe a disturbance of the microbial community composition during viral infections that does not return to baseline after the acute viral infection resolves. Moreover, we show that *Pseudomonas* relative abundance is significantly increased in the airways of pwCF during viral infection and that increased concentrations of antiviral cytokines – such as interferon (IFN)-*λ*1 – are associated with increased *Pseudomonas* abundance. These findings offer evidence that the progression of chronic *Pseudomonas* infections in pwCF are influenced by acute respiratory viral infections and the subsequent antiviral response in the airways. This study furthers our understanding of viral-bacterial coinfection in the context of CF.

## INTRODUCTION

People with cystic fibrosis (pwCF) commonly experience chronic bacterial infections of the airways that take hold in adolescence and persist throughout their lifetime^1–3^. Cystic fibrosis (CF) is caused by a mutation in the CF transmembrane conductance regulator (CFTR) gene which encodes for a chloride ion transport channel. Incorrect ion homeostasis at mucosal epithelial sites throughout the body lead to thick, dehydrated mucus which leads to damage to multiple organ systems^4^. This is particularly problematic in the airways where thickened mucus is not able to be sufficiently cleared and so forms an ideal environment for inhaled microbes to grow. Chronic infections lead to chronic inflammation and damage to the airways, leading to reduced lung function and increased morbidity and mortality^5,6,1,7–9^.

*Pseudomonas aeruginosa* is an important pathogen in the CF airways, associated with increased morbidity and mortality in pwCF. It is frequently associated with water sources in proximity to human activity and, as such, is a common nosocomial opportunistic pathogen^10^. Aggressive eradication treatments are standard in pediatric treatment regimens, but over time antibiotic treatment becomes less and less effective until patients are considered chronically infected^11–14^. The cause of the switch from intermittent to chronic *P. aeruginosa* infection has long been a subject of investigation in CF research. It has been previously observed that the transition to chronic infection occurs more frequently during winter months, correlating with respiratory virus season in the northern hemisphere^15^. In pediatric patients, it has also been shown that 83% of new *P. aeruginosa* infections occur after a preceding viral infection^16^. In adults, more than 30% of exacerbations occurred when the patient tested positive for a respiratory viral infection^17^. pwCF do not contract respiratory viruses at rates greater than the general population but experience substantially worse outcomes^18,19^. Viral infections can trigger pulmonary exacerbations – acute worsening of pulmonary symptoms and lung function – which are associated with intravenous antibiotic usage and longer hospital stays^20–24^. Following an exacerbation lung function often does not return to pre-exacerbation levels, meaning lung function is gradually decreased until respiratory failure or need for a lung transplant^25^. There is little data about the incidence or severity of respiratory viral infections in the age of highly effective modulator therapies (HEMT) which were first approved by the FDA in 2019. More than 90% of pwCF^1^ are eligible for HEMT, although it is not universally well tolerated^26^. After starting treatment pwCF experience significant increases in lung function, lessening of symptoms, and decreased pathogen abundance^27–30^. Although, pathogens are not eradicated in the airways post-HEMT^29,30^ and there are many pwCF for whom HEMT are not available or not approved, so chronic infection remains an important area of research in CF.

We have previously shown that *P. aeruginosa* biofilm growth is increased on CF bronchial epithelial cells secondary to acute respiratory viral infection^31^ and the subsequent antiviral interferon response^31,32^. During viral infection, antiviral type I and III interferons (IFN) are secreted and signal through receptors on the epithelial cells which triggers metabolic reprogramming^33,34^ and nutritional immunity dysfunction, which changes the secreted milieu in which *P. aeruginosa* resides. Specifically, there are increases iron bioavailability^33^ and secreted metabolites, including L-lactate^32^ which can all promote biofilm production by *P. aeruginosa*. The biofilm mode of growth, in which bacterial cells grow in a multicellular community encased in a protective extracellular matrix, is a hallmark of chronic infection in pwCF^35–39^. Biofilms are recalcitrant to antibiotic treatment and increase resistance of *P. aeruginosa* to killing by host immune cells. Further to this, we have also shown that increased iron concentration in CF sinuses is associated with decreased alpha diversity of the bacterial community^33^ indicating that post-viral nutritional and metabolic reprogramming of airway cells may drive dominance of *P. aeruginosa* in the community.

We hypothesize that pwCF experience increases in *P. aeruginosa* abundance in the airway with viral infection, potentially leading to dominance of the microbial community and chronic infection. In this study, we aim to identify changes in the microbiota of the CF airway during and after viral infection, as well as impacts of host antiviral response on bacterial pathogens in the airway.

## RESULTS

### Study cohort

Participants were enrolled in an observational study at the University of Pittsburgh Medical Center Adult CF Sinus Clinic following an IRB-approved protocol (STUDY19100149). All participants provided informed consent upon enrollment into the study. Samples were collected from the upper and lower airways in the form of sinus swabs for microbiota analysis and viral reverse transcriptase (RT)-PCR panel (**figure S1**), sinus aspirates for cytokine biomarker analysis, and expectorated sputum and microbiota and viral RT-PCR panel (**figure S1**). Sinus samples were collected under direct endoscopic observation from the right maxillary sinus and swabs were protected by a sheath during sampling to protect from contamination by the nares. Where possible, all sample types were collected during every visit. Samples were collected from 38 adults with CF and concomitant chronic rhinosinusitis (CRS) approximately every 3 months, with extra sample visits during pulmonary exacerbations, over the course of 42 months from 2017 – 2021. Participants enrolled in the study had previously undergone functional endoscopic surgery (FESS) to alleviate CRS symptoms. Microbiota profiles from sinus and sputum that did not meet quality thresholds were excluded from this analysis (**figure 1**). Samples from a total of 36 participants were included in final analysis (**table 1**). We collected a total of 202 samples of which 128 were sinus and 74 were sputum samples. Of these, 24 sinus samples (19%) and 10 sputum samples (14%) were positive for a viral pathogen by RT-PCR panel testing (**table 1**). Viral RT-PCR panel tested for acute respiratory viruses as detailed in Ali *et al.* (2011)^40^ and ten different viruses were identified in our samples: human metapneumovirus, human rhinovirus (hRV), influenza A, parainfluenza viruses 1 – 4, respiratory syncytial virus (RSV), and seasonal coronaviruses 229E and OC43 (**figure S1**). The most frequently identified virus in both sinus and sputum samples was hRV at 54% and 60%, respectively.

**table 1.**

|  |  |
| --- | --- |
| Total subjects in this study = 36 |  |
| Median age at enrollment (years; range) | 29.95 (21.21 – 49.72) |
| Male (%) | 16 (44%) |
| Race (%) |  |
| White | 35 (97%) |
| Other | 1 (3%) |
| Cystic fibrosis transmembrane conductance regulator (CFTR) genotype (%) |  |
| ΔF508 homozygous | 19 (53%) |
| ΔF508 heterozygous | 15 (42%) |
| Other | 2 (5%) |
| Allergic rhinitis (%) | 14 (39%) |
| Viral positive samples (%) |  |
| Sinus | 24 (19%) |
| Sputum | 10 (14%) |

**figure 1.**
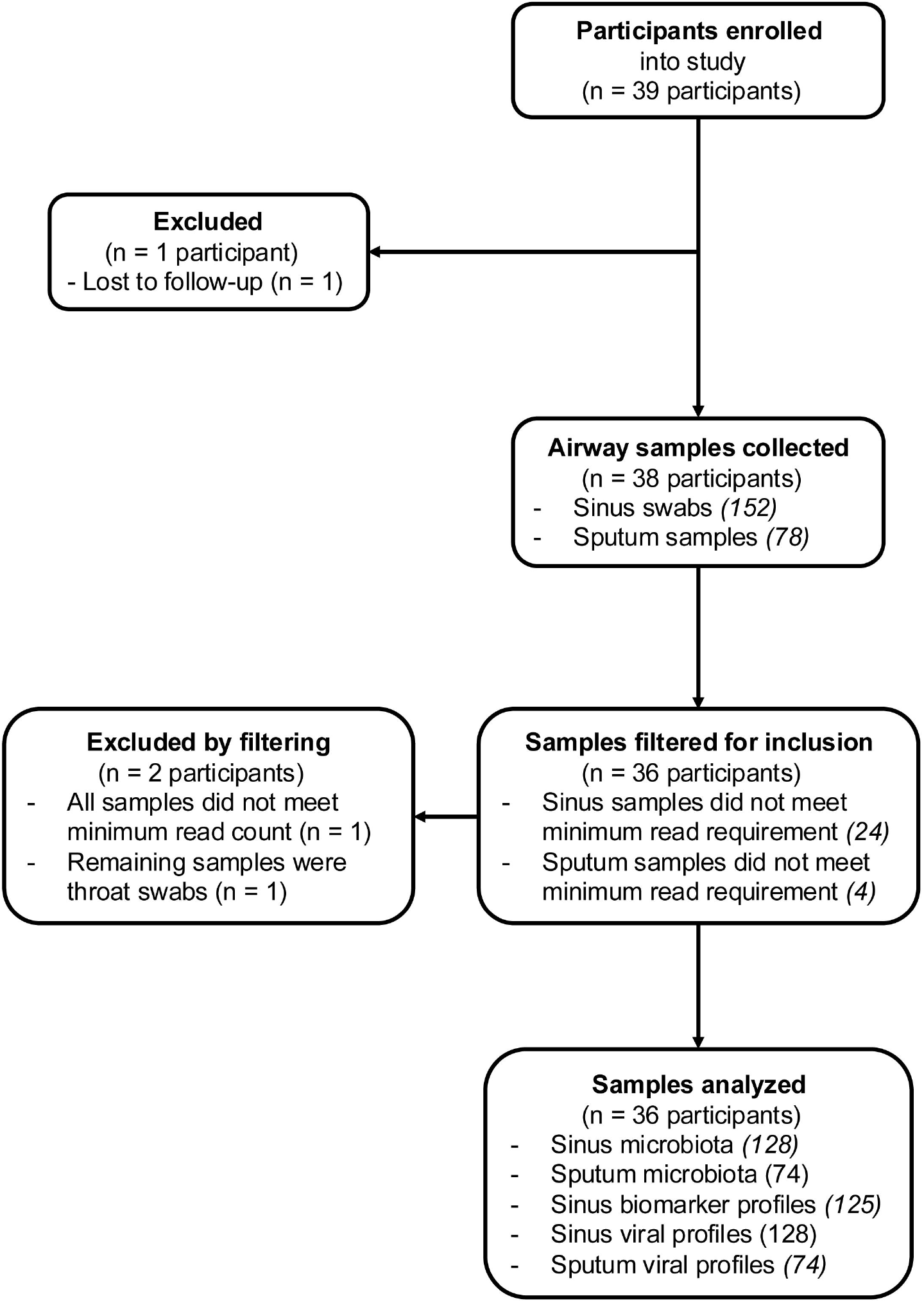

### Airway microbiota are perturbed during respiratory viral infection

To assess if respiratory viral infections lead to changes in the airway microbiota of pwCF, we grouped samples based on their temporal proximity to a positive viral sample (<u>pre</u>-virus, <u>active</u> viral infection, and <u>post</u>-virus). Per participant centroids were generated for all samples in the pre-virus subset. Distance to centroid, a measure of variation between communities, was calculated for all samples by participant (i.e. each sample distance was only measured to the pre-virus centroid for the participant who provided the sample). Distance to centroid was compared between viral states. Distance to centroid was low (median = 0.23) when comparing pre-virus samples to their pre-virus centroid (**figure 2**). This reflects within-participant variation in bacterial community composition which has been previously noted in pwCF^30,41,42^. There is a significant increase in distance to pre-virus centroid for samples collected during viral infection (median = 0.75, P < 0.0001) indicating substantial variation in the bacterial community composition during viral infection. The increase in distance to pre-virus centroid for samples collected post-viral infection is less than during viral infection (median = 0.37) but is still significantly different from the pre-virus samples (P = 0.01), indicating that the composition of airway microbiota is not entirely restored to baseline following viral infection.

**figure 2.**
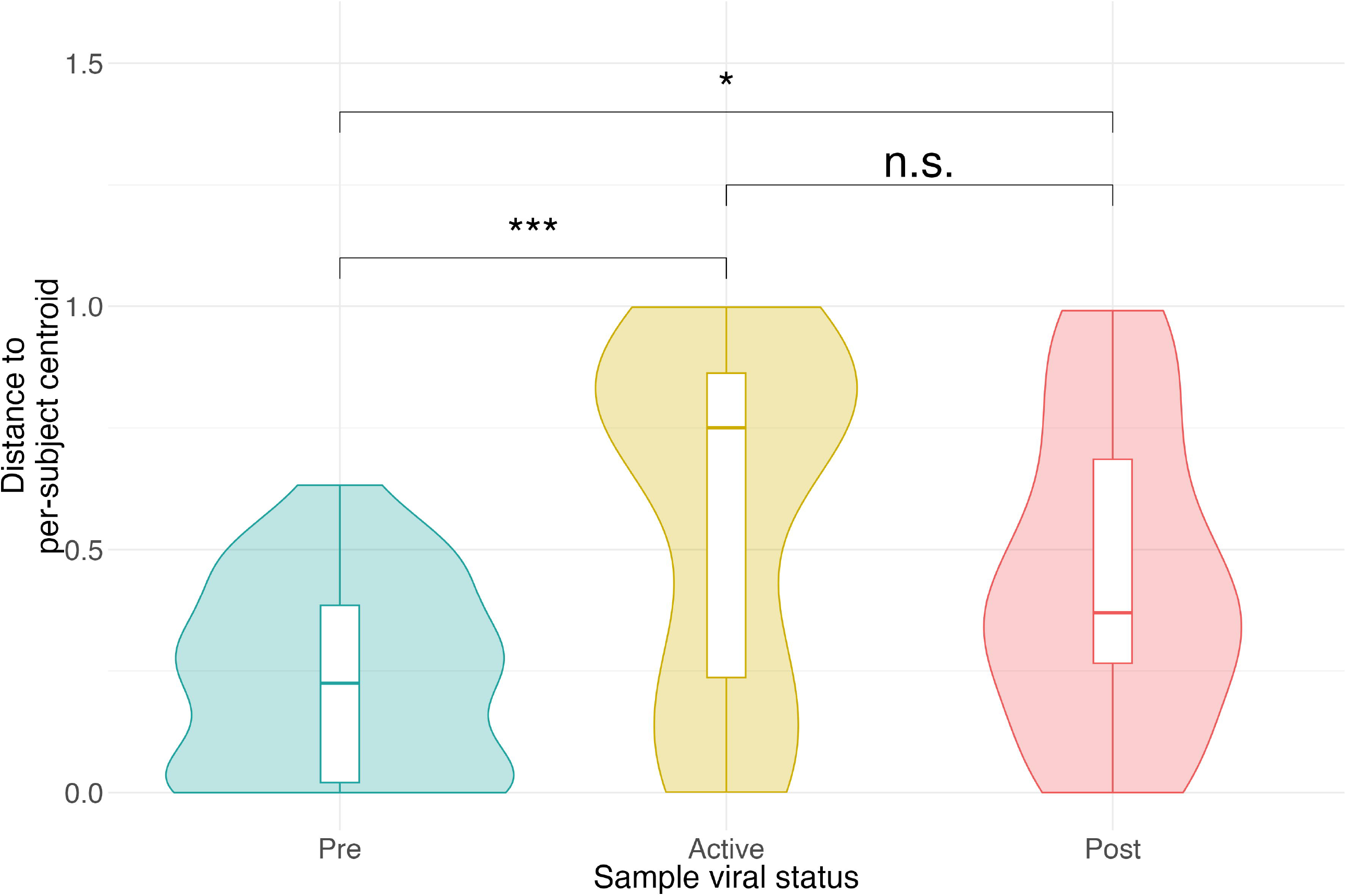

### *Pseudomonas* abundance is increased during viral infection

To identify the taxa responsible for shifts in microbial community composition during viral infection we first sought to identify core and satellite taxa of the adult CF airways using previously published methodology^43^ (**figure 3A**). Core taxa were defined as those with prevalence across all samples >75%. We identified three core taxa (*Staphylococcus, Pseudomonas*, and *Streptococcus*) and 96 satellite taxa. Interestingly, the core taxa were not evenly distributed throughout the airway. *Streptococcus* was observed more frequently in sputum samples than in the sinus, and often in the sputum of participants who did not have *Streptococcus* identified in their paired sinus samples (**figure 3B**).

**figure 3.**
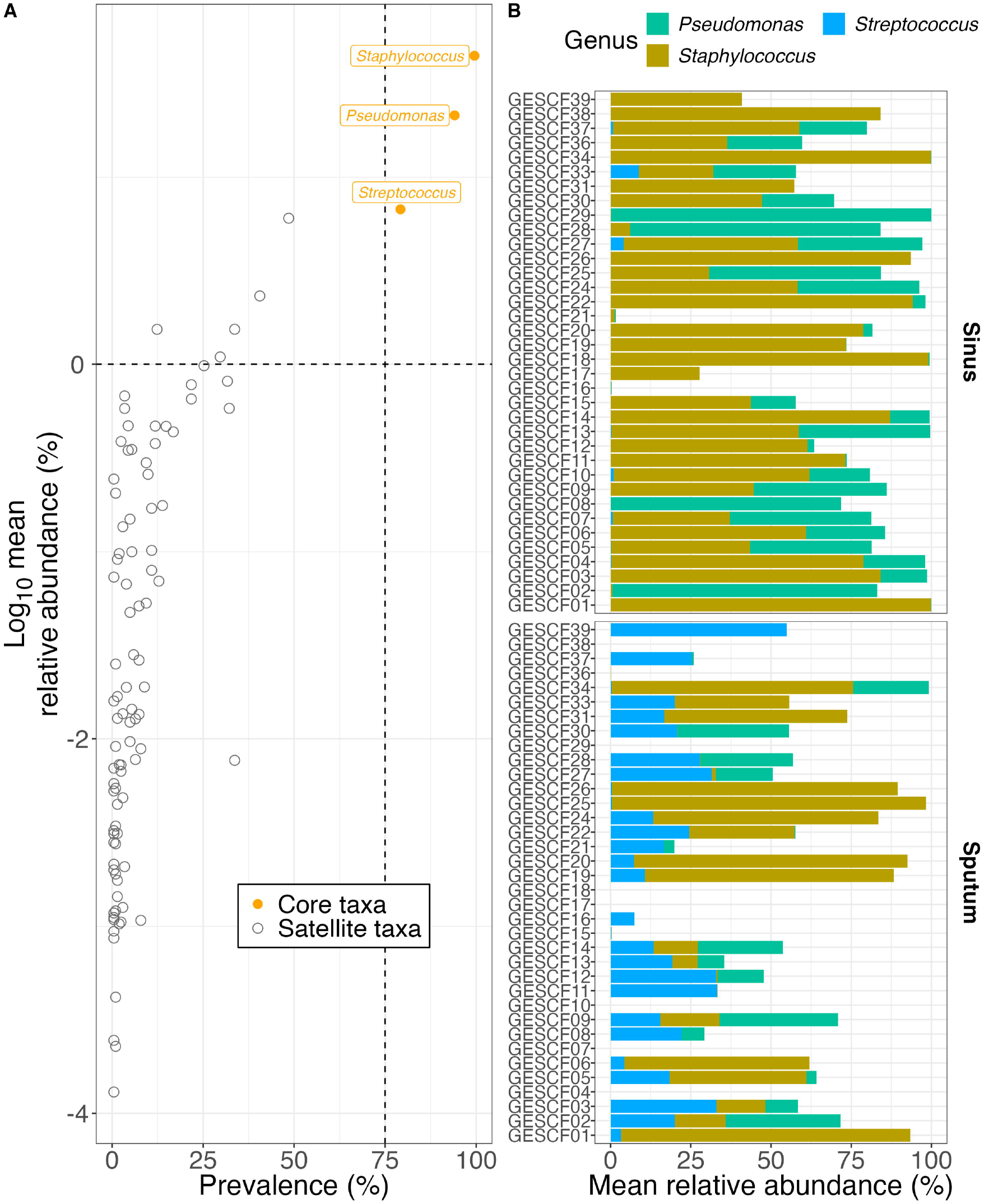

By examining changes to core taxa relative abundance over time and in relation to positive viral samples, we observe alterations to relative abundance for all three genera during viral infection (**figure 4A**). When comparing samples based on viral status, there was a significant increase in *Pseudomonas* relative abundance (P = 0.029) in positive viral samples compared to negative ones (**figure 4B**). We observed a trend towards decreased *Staphylococcus* relative abundance in viral positive samples. The bimodal distribution of relative abundance of *Pseudomonas* and *Staphylococcus* during viral infection is representative of participants whose samples were consistently dominated by *Staphylococcus* and did not experience a microbiota perturbation during viral infection. These data suggest that acute respiratory viral infection results in increased *Pseudomonas* in the airways of pwCF^32–34^ and may contribute to *Pseudomonas*’ path to dominance in the airways over the course of chronic infection.

**figure 4.**
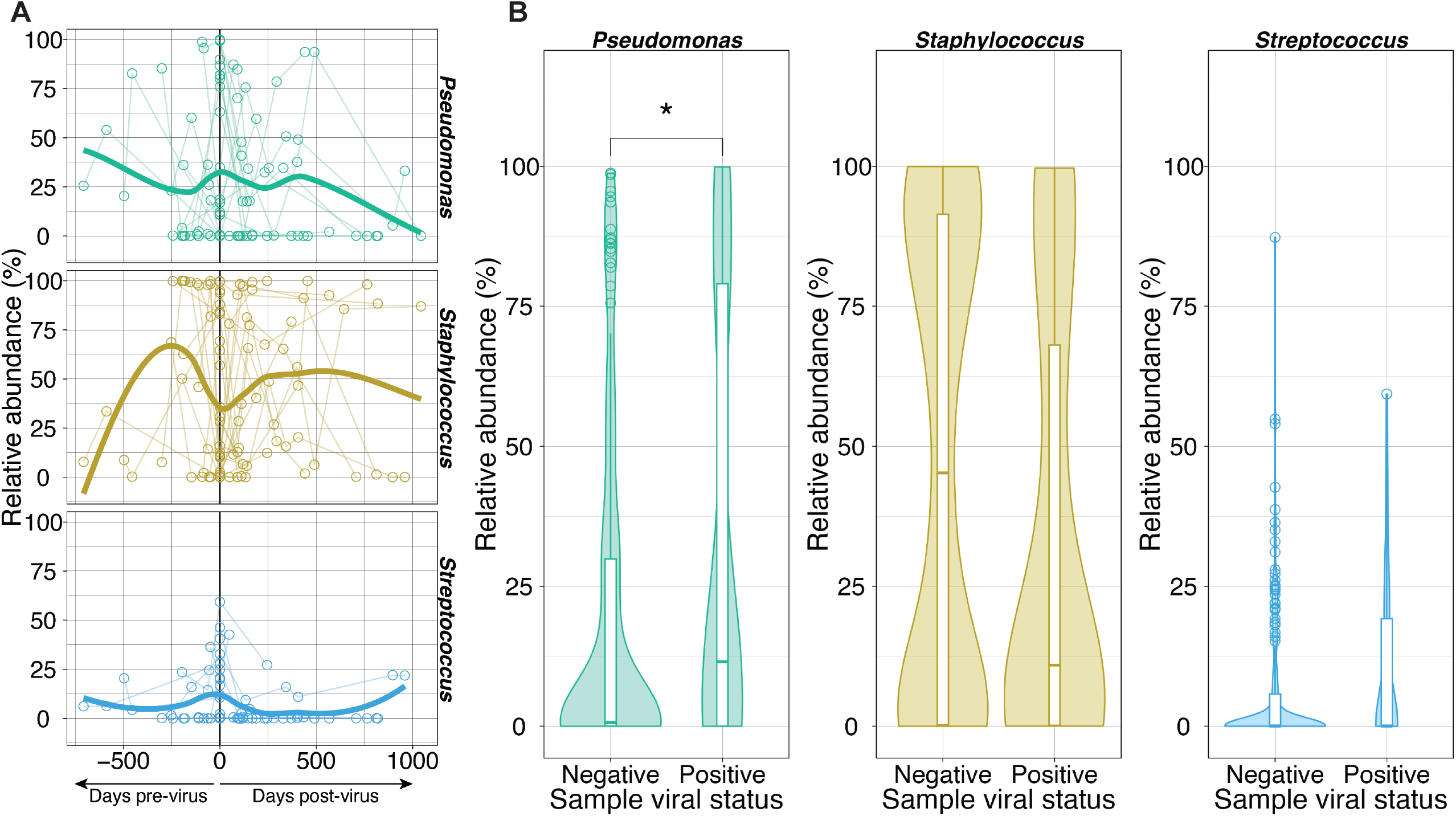

### Respiratory viral infection significantly increases antiviral and anti-inflammatory cytokines in the CF airway

A panel of 37 cytokines were assessed from sinus swabs (**figure S2, table S1**) and we observed that seven of these had significantly increased concentrations in viral positive samples vs. viral negative samples (**figure 5, table S2**). We observed a slight but significant increase in interferon (IFN) gamma (IFNg, IFN-*γ*) in viral positive samples (P = 0.03) as well as an increase in three interleukin (IL) cytokines; IL-8 (P = 0.047), IL-10 (P = 0.029), and IL-22 (P = 0.029). Increases were also observed in B cell-activating factor (BAFF) (P = 0.034), glycoprotein 130 (gp130) (P = 0.061), and soluble tumor necrosis factor receptor 1 (sTNFR1) (P = 0.045) which play roles in adaptive immunity, promotion of inflammation, and suppression of immune response, respectively. These data indicate that, although the CF airway is persistently and chronically inflamed, acute viral infections play a role in modulating the inflammatory environment in which airway microbiota reside.

**figure 5.**
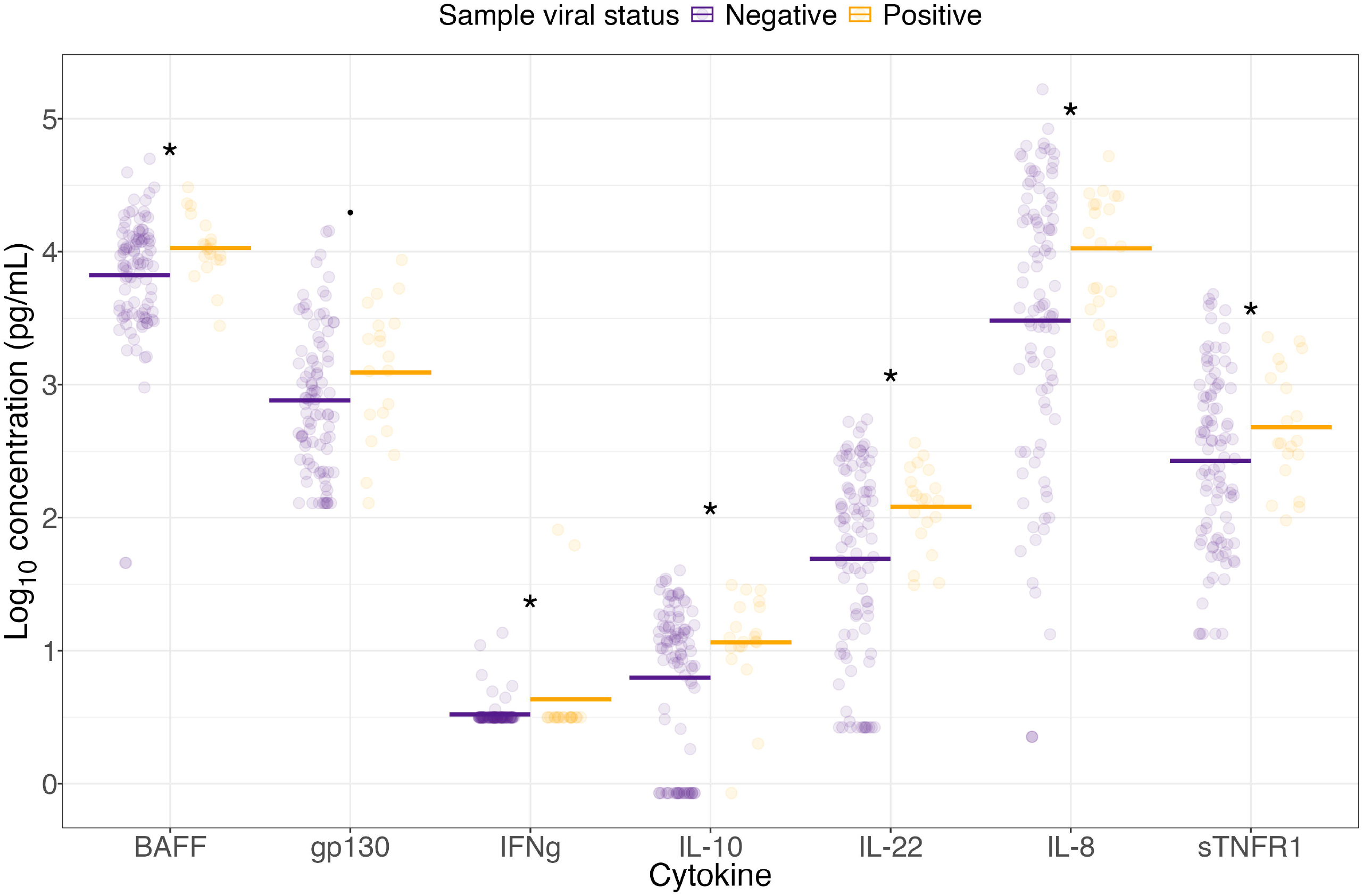

### Specific cytokines are associated with increased *Pseudomonas* abundance during viral infection

Since our previous studies showed increased *P. aeruginosa* biofilm growth in response to antiviral cytokine signaling during acute viral infections, we next assessed if changes in cytokine levels during viral infection correlated with relative abundance changes in *Pseudomonas*. Linear mixed effect modeling in which cytokine concentration and sample viral status were input as an interaction term revealed significant associations between concentrations of seven cytokines and *Pseudomonas* relative abundance during viral infection (**figure 6**). IL-8 (P = 0.0158) and IL-22 (P = 0.070) are important cytokines in the human antiviral response and are associated with increased *Pseudomonas* relative abundance. IL-2 (P = 0.069) plays a role in T cell activation and is likely increased as part of the antiviral response in the airway. Chitinase-3 like-protein 1 (CHI3L1) (P = 0.029) and IL-35 (P = 0.031) are both mediators of a pro-inflammatory response via induction of other cytokines. sTNFR2 (P = 0.585) acts to bind TNF cytokines and suppress inflammatory response. Importantly, increased levels of IL-29 (IFN-*λ*1) during viral infection were shown to be significantly associated with higher *Pseudomonas* relative abundance (P = 0.0172). There are many host factors involved in antiviral immune response and these data suggest that increases in *Pseudomonas* biofilm observed *in vitro* when stimulated with respiratory viruses and IFN^32^ may also occur *in vivo*. As has been shown for acute viral-bacterial co-infections^44^, we observe the that antiviral IFN response is also associated with worsening secondary bacterial infections in a chronic infection setting.

**figure 6.**
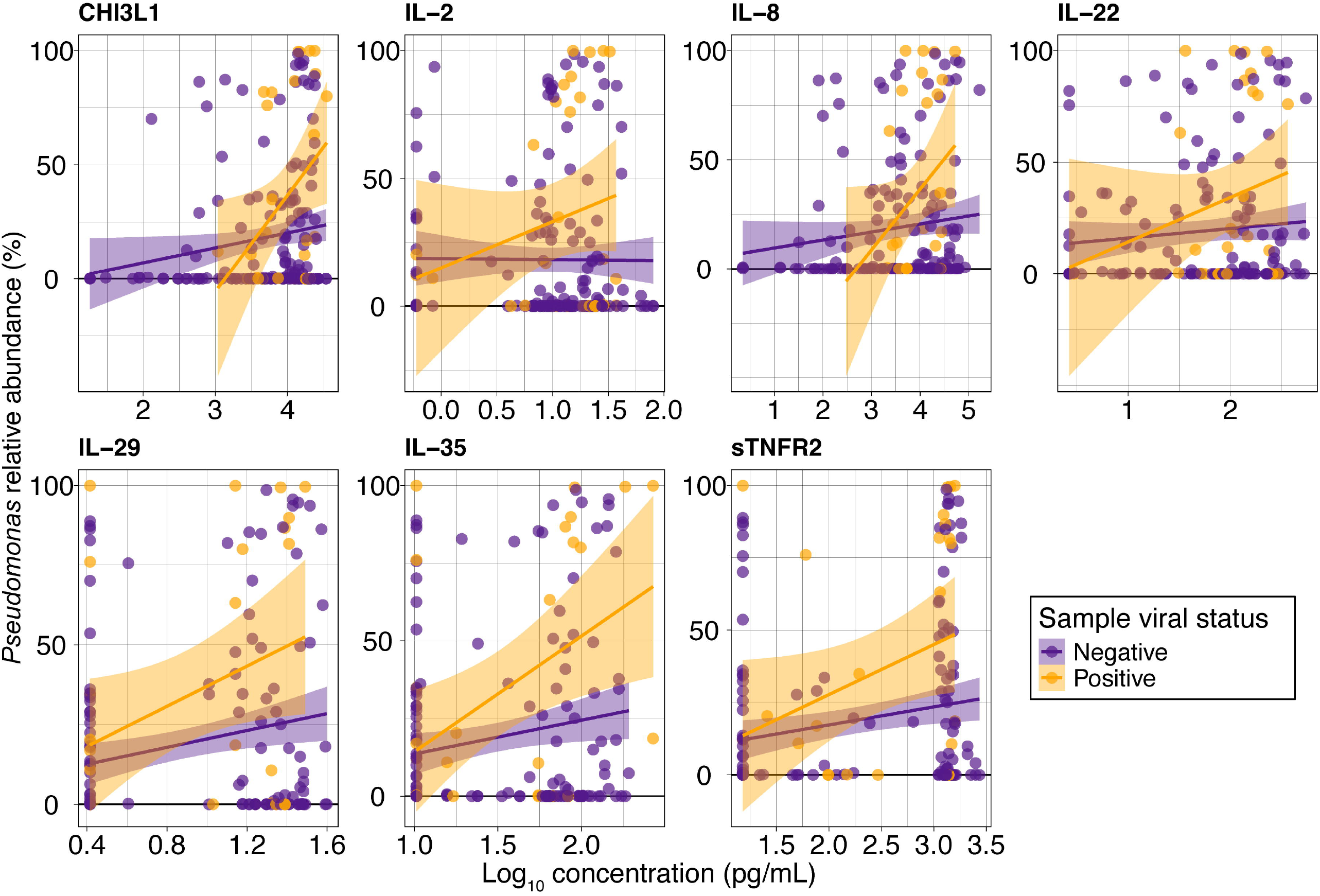

## DISCUSSION

This study sought to identify changes in the microbiota of the CF airway induced during respiratory viral infection that shape the outcomes of chronic bacterial infections in pwCF. There is significant perturbation of the CF airway microbiota during viral infection that does not return to baseline after viral resolution. We observed a significant increase in the relative abundance of *Pseudomonas* in the airways of pwCF during respiratory viral infection. Furthermore, we report associations between antiviral cytokine response and the increase in *Pseudomonas* abundance during co-infection.

Respiratory viral infections have been shown in numerous studies to play a role in exacerbations and worsened symptoms in pwCF^20,18,22,17^ but little is known about alterations to the resident airway microbiota during viral infection. Here, we show that the community composition of the CF airways is significantly altered during viral infection. We observed that distance to centroid is significantly increased both during and post-viral infection when compared to pre-infection samples (**figure 2**). Increased distance to centroid during viral infection indicates that bacterial community composition is altered, either by drastic changes to relative abundance of existing community members or by the loss or gain of community members. We observed a smaller but significant increase in distance to centroid when comparing post-virus samples to their pre-virus centroids. This likely represents a shift in community composition toward the pre-virus state, but clearance of the virus does not entirely restore the community as it previously existed. These data support the hypothesis that viral infections play a role in remodeling the bacterial community structure in the CF airways and drive pathogens towards chronic infection and dominance of the microbiota.

The airway microbiota of pwCF tend to have low alpha diversity (few observed species and communities dominated by one or two taxa) but often varies widely between and within participants^42,30^. It can be challenging to identify patterns of change in longitudinal datasets when variation within participants is high. To better understand the role of individual genera in community restructuring during viral infection, we identified the core taxa present in our samples. We identified core taxa from the CF airways as *Staphylococcus, Pseudomonas*, and *Streptococcus. Streptococcus* was shown to be more prevalent in the lower airway than in the sinus (**figure 3**). We observed significantly increased *Pseudomonvas* relative abundance and a substantial reduction in *Staphylococcus* relative abundance during viral infection (**figure 4B**). In many participants, *Pseudomonas* relative abundance decreased post-viral infection (**figure 4A**) indicating that active viral infection is a key driver of the increases in relative abundance. We have previously shown that viral infection of airway epithelial cells increases iron bioavailability^31,33^ and that *in vivo* iron levels are correlated with increased transcription of a type VI secretion system toxic effector (TseT) and decreased alpha diversity in the sinuses of pwCF^33^. Taken alongside these previous studies, our current data provides evidence for a link between acute respiratory viral infection and increase abundance of *Pseudomonas* in the CF airways.

We sought to confirm that acute viral infections of the CF airway induce an antiviral response and we have demonstrated that several important antiviral, pro-, and anti-inflammatory cytokines are present at higher concentrations in the airways of pwCF during viral infection (**figure 5**). Inflammation of the CF airway is chronic and inflammatory biomarkers quantified in the airways of pwCF are significantly higher than in healthy participants ^45–47^. The small number of cytokines with observed increases in concentration during viral infection is likely due to the chronic nature of CF airway inflammation, where inflammatory biomarkers are consistently present at elevated concentrations.

We observed that several cytokines, with varying roles in the immune response, were associated with increased *Pseudomonas* relative abundance during viral infection (**figure 6**). Pro-inflammatory cytokines chitinase-3 like-1 (CHI3L1), IL-8, and IL-35 were all found to be positively associated with increased *Pseudomonas* during viral infection, as were IL-22 and IL-29 (IFN-*λ*1) (antiviral cytokines), IL-2 (a T cell regulator), and sTNFR2 (an anti-inflammatory receptor). The variation among the roles and superfamilies of these cytokines implicated in increased *Pseudomonas* relative abundance raises questions about the mechanisms by which *Pseudomonas* spp. can sense and interact with the host environment to leverage a situational advantage. Approximately 8% of the *P. aeruginosa* genome encodes for regulatory genes^48^ which allows for rapid and effective responses to environmental stimuli, helping *P. aeruginosa* thrive in environments hostile to many other bacterial species. Viral infection of airway epithelial cells causes IFN secretion and subsequent metabolic reprogramming as part of the antiviral response^32^. IFNs are secreted as part of the innate immune response to viral infection and we previously demonstrated that human CF bronchial epithelial cells infected with respiratory syncytial virus (RSV) undergo a metabolic shift towards aerobic glycolysis which leads to increased apical secretion of L-lactate. In response to increased L-lactate, *P. aeruginosa* exhibits increased biofilm formation^32^ indicating that viral infection can drive secondary bacterial infections and expansion of existing *P. aeruginosa* populations in the CF airways. Our findings in this study provide compelling evidence that our *in vitro* observations^31–33^ of the effects of viral infection on *P. aeruginosa* are accurately recapitulating microbial interactions in the CF airway.

There are some limitations to our study. Firstly, we were unable to utilize shotgun metagenomics in this study due to low bacterial biomass and high host burden in our samples, particularly sinus swabs. In preliminary experiments, host reads made up >99% of the total reads in most samples. When unable to reconstruct *de novo* bacterial genomes from the remaining reads, we were only able to assign taxonomy to reads by using a single copy gene database^49^. Using 16S rRNA amplicon sequencing produced more consistent results but prohibited identification of taxa in our dataset beyond genus level and constrained our analysis by the compositionality of relative abundance data. These data were collected only from pwCF with symptomatic CRS and history of FESS which may impact the applicability of the results to a broader CF population. The data in this study also do not take into account the effects of HEMT on the prevalence of viral infections or their effects on the airway microbiota of pwCF. There have been many early studies published examining the symptomatic improvements and reductions in pathogen burden in pwCF using HEMT7/8/25 7:58:00 AM but we are yet to see data from long-term studies that may reveal the changing landscape of infection and disease management in an aging population with CF.

## CONCLUSION

We have shown that *Pseudomonas* relative abundance is increased in the airways of pwCF during viral infection and that this perturbation to the microbiota is not resolved once the viral infection is cleared. Increased *Pseudomonas* abundance is correlated with increased antiviral cytokines indicating that repeated viral infections drive proliferation of existing *Pseudomonas* populations and may lead to chronic infections experienced by pwCF as they age. These data provide compelling evidence that the progression of chronic *Pseudomonas* infections in pwCF are influenced by repeated acute respiratory viral infections.

## METHODS

### Sample collection

We performed a prospective longitudinal study of adults attending a CF-focused otolaryngology clinic at the University of Pittsburgh Medical Center (Pittsburgh, PA) following an IRB-approved protocol (STUDY19100149). All participants provided informed consent upon enrollment into the study. Thirty-nine adults were enrolled; one was lost to follow-up and two were omitted from this analysis due to samples not meeting quality criteria (**figure 1**). All participants had diagnosed CF and symptomatic CRS and had previously undergone FESS to alleviate CRS symptoms. Demographic data were collected upon enrollment and airway comorbidities such as asthma and allergic rhinitis were determined by retrospective review of pulmonology and otolaryngology notes. Participants attended scheduled quarterly visits to the clinic as well as unscheduled visits during pulmonary exacerbations. Sinus samples were collected under direct endoscopic observation from the right maxillary sinus. Swabs for 16S rRNA amplicon sequencing and viral panel screening (nylon flocked swab; Puritan Medical Products, Guildford, ME) were collected by insertion into the sinus through a sterile sheath to ensure no contact with the nares during sampling. The sinus was rinsed with 5 mL sterile saline and aspirated into a sterile trap for cytokine biomarker analysis. Expectorated sputum was collected in a sterile specimen container (Covidien general purpose specimen container, Fisher Scientific, Waltham, MA). All samples were transported on wet ice from the clinic to the laboratory (University of Pittsburgh, Pittsburgh, PA) before storage at -80°C.

### DNA extraction

DNA was extracted from samples sorted by participant and samples type (e.g. sinus swab or sputum) and extracted in lots to reduce batch effects for intra-participant comparisons. An extraction control sample was included in each batch and consisted of an unused sterile flocked swab or sterile distilled water that was subjected to the complete DNA extraction and purification protocol. The protocol “Benzonase 2” was carried out as described in Nelson *et al.*^50^ with an alteration to the hypertonic lysis step for sinus swabs: host cells are more readily accessible on a nylon flocked swab than in sputum and so sinus swabs were incubated in sterile distilled water at room temperature for 15 min.

### Viral panel RT-PCR

Sinus swab samples were processed to assess presence of viral pathogens by real-time RT-PCR using previously published primers and methodology ^40^. Viral RNA was extracted from swabs and stored at -80°C until use. Single-plex RT-PCR reactions were carried out for all viruses in the panel and suitable negative and positive controls were included in each assay. Fourteen viruses were included in the panel: respiratory syncytial virus (RSV), influenza A and B, human metapneumovirus, human rhinovirus (hRV), parainfluenza 1, 2, 3 and 4 (PIV1, PIV2, PIV3, PIV4), adenovirus and human coronaviruses Netherlands (NL-63), OC43, 229E, and SARS-CoV-2 coronavirus.

### Cytokine biomarker analysis

Cytokine concentrations were quantified in sinus aspirate samples using a Luminex MAGPIX system (Luminex, Austin, TX) with a Bio-Plex Pro Human Inflammation Panel 1, 37-Plex #171AL001M kit (Bio-Rad, Hercules, CA) following manufacturer’s guidelines.

### 16S rRNA amplicon sequencing

Extracted and purified DNA from samples and controls was sent to DNA Services at the University of Illinois at Urbana-Champaign, IL for amplification, library preparation, and sequencing. Amplicons were generated from the 16S V4 variable region (515f – 806r). Amplicon libraries were quantified by qPCR and sequenced on a NovaSeq flowcell for 251 cycles from each end of the fragment using a NovaSeq 500-cycle sequencing kit v1.5 (Illumina, San Diego, CA). Generated fastq files were demultiplexed using bcl2fastq v2.20 Conversion Software (Illumina, San Diego, CA).

### 16S rRNA sequence analysis

All code used for this analysis can be found in Supplemental data file S1. Briefly, 16S V4 amplicon data were important into QIIME2 v2021.11^51^ using the “EMPPairedEndSequences” option. Sequences were demultiplexed using no Golay error correction and denoised using the DADA2^52^ plug-in. Chimeric sequences were removed using the VSEARCH^53^ plug-in. Samples were rarefied to an optimal depth of 11,500 to maintain the maximum number of observed features before classifying taxa using the SILVA 138.1 rRNA database^54^. Feature data were converted to a BIOM table^55^ and downloaded as a tab-separated text document for further analysis in R v4.2.1.

Data were imported to phyloseq v1.40.0^56^ and amplicon sequence variants (ASVs) were decontaminated using frequency and prevalence data in samples and controls using the package decontam v1.16.0^57^. Non-bacterial taxa were filtered from the data set in phyloseq and duplicate genera were agglomerated using the function “tax_glom” and specifying genus as the taxonomic rank. CHAO1, observed, Shannon, and Simpson diversity indices were estimated using phyloseq. Taxa labeled “unassigned” at the genus level were identified and their V4 variable region amplicons extracted from the representative sequence file. Unassigned 16S sequences were queried using NCBI BLAST and sequences with a >99% similarity to a given organism were manually assigned to that genus. Relative abundance per ASV per sample was calculated and these values were used for further analysis.

Prior to beta diversity analysis, samples were filtered to a total minimum read count of 1045; samples with fewer reads were omitted from further analysis (**figure 1**). Beta diversity analysis was carried out using vegan v2.6.4^58^. Morisita-Horn index was calculated from relative abundance data. Centroids and distance to centroid were calculated from Morisita-Horn index matrix using vegan and a custom function written by Jari Oksanen taken from the vegan package github page.

Differences in relative abundance and relationships between abundance and cytokine concentrations were tested by application of mixed effect linear models using lme4 v1.1.34^59,60^ in which participant ID was set as a random effect. Statistically significant differences between linear models were tested using ANOVA with post-hoc Tukey test where necessary.

## ACKNOWLEDGEMENTS

The authors would like to thank all participants in this study for their time and support of CF research. Thanks also go to Ryan Little, MD for his advice and feedback in preparation of this manuscript.

This work was supported by the National Institutes of Health (NIH) grant R33HL137077 and a GILEAD Investigator Sponsored Research award to S.E.L., V.S.C., and J.M.B.; and NIH grants R01HL123771, R01HL142587, and P30DK072506, and Cystic Fibrosis Foundation (CFF) grants BOMBER18G0 and RDP BOMBER19R0 to J.M.B., CFF Carol Basbaum Memorial Research Fellowship (ARMBRU19F0), CFF Postdoc-to-Faculty Transition Award (ARMBRU22F5), and the NIH grant T32HL129949 to C.R.A..

