## Supplementary figures and images for "Respiratory viral infection is associated with increased *Pseudomonas* abundance in cystic fibrosis airways"

### Figure S1

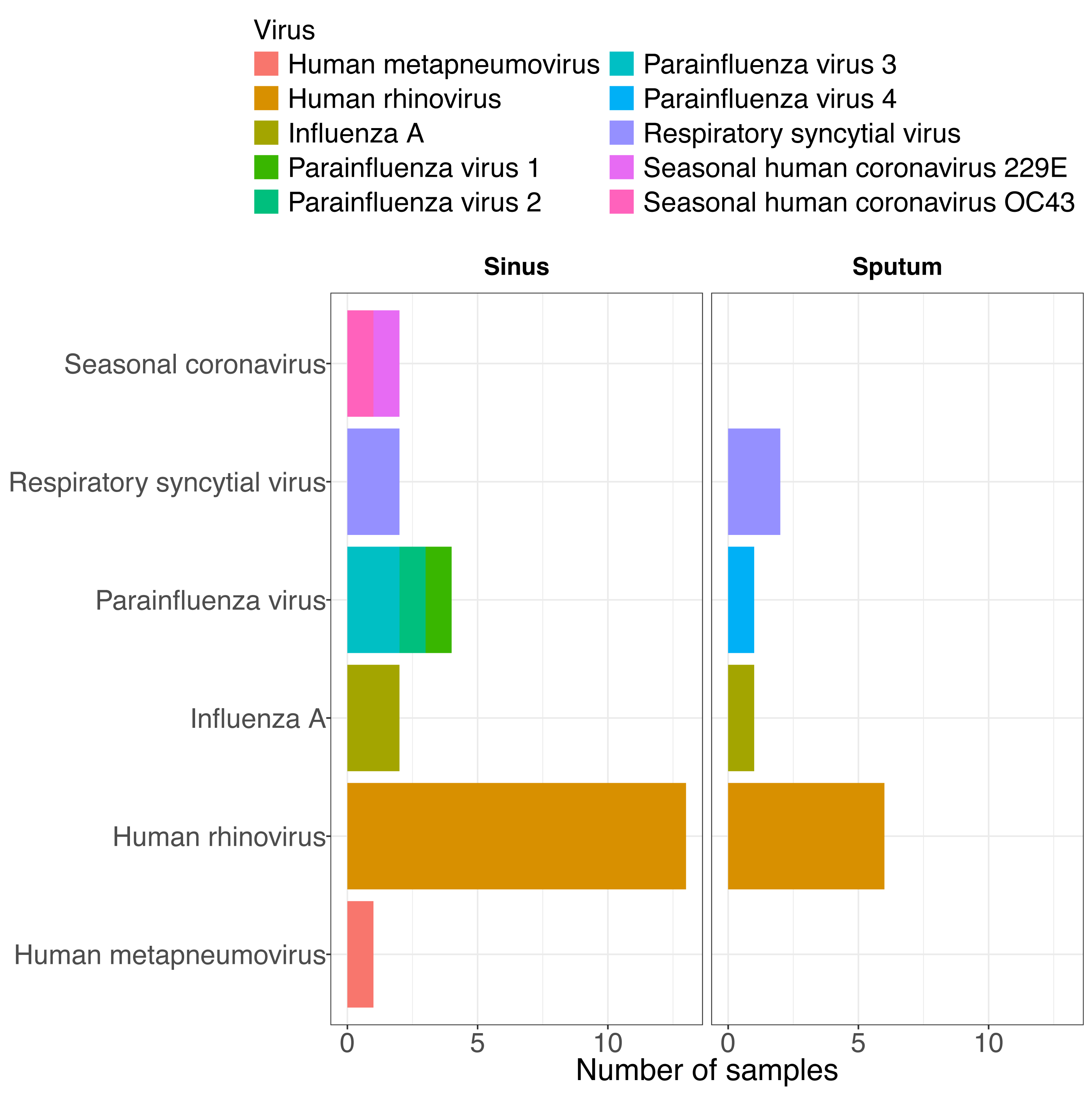

### Figure S2

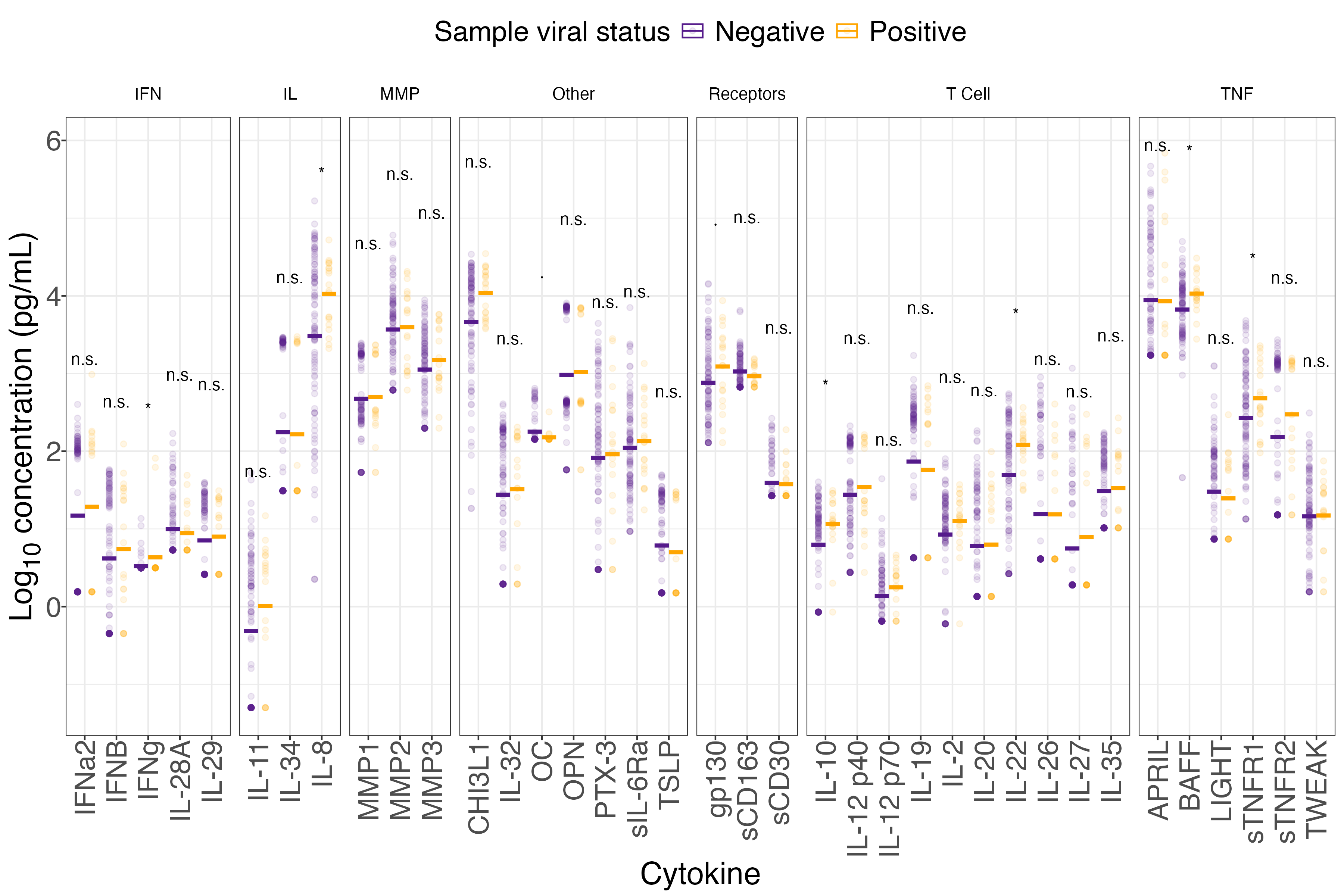
